# From Visual Exploration to Storytelling and Back Again

**DOI:** 10.1101/049585

**Authors:** Samuel Gratzl, Alexander Lex, Nils Gehlenborg, Nicola Cosgrove, Marc Streit

## Abstract

A major challenge of data-driven biomedical research lies in the collection and representation of data provenance information to ensure reproducibility of findings. In order to communicate and reproduce multi-step analysis workflows executed on datasets that contain data for dozens or hundreds of samples, it is crucial to be able to visualize the provenance graph at different levels of aggregation. Most existing approaches are based on node-link diagrams, which do not scale to the complexity of typical data provenance graphs. In our proposed approach we reduce the complexity of the graph using hierarchical and motif-based aggregation. Based on user action and graph attributes a modular degree-of-interest (DoI) function is applied to expand parts of the graph that are relevant to the user. This interest-driven adaptive provenance visualization approach allows users to review and communicate complex multi-step analyses, which can be based on hundreds of files that are processed by numerous workflows. We integrate our approach into an analysis platform that captures extensive data provenance information and demonstrate its effectiveness by means of a biomedical usage scenario.

## 1 Introduction

Scientific progress is driven by discoveries based on observations. Accurate and efficient documentation and presentation of how discoveries were made is essential, since the scientific method requires that findings are reproducible. The process from making a discovery in a visualization tool to communicating it to an audience is typically a process that does not allow users to switch back from presentation to exploration, as illustrated in Figure 2.

**Figure 1:**
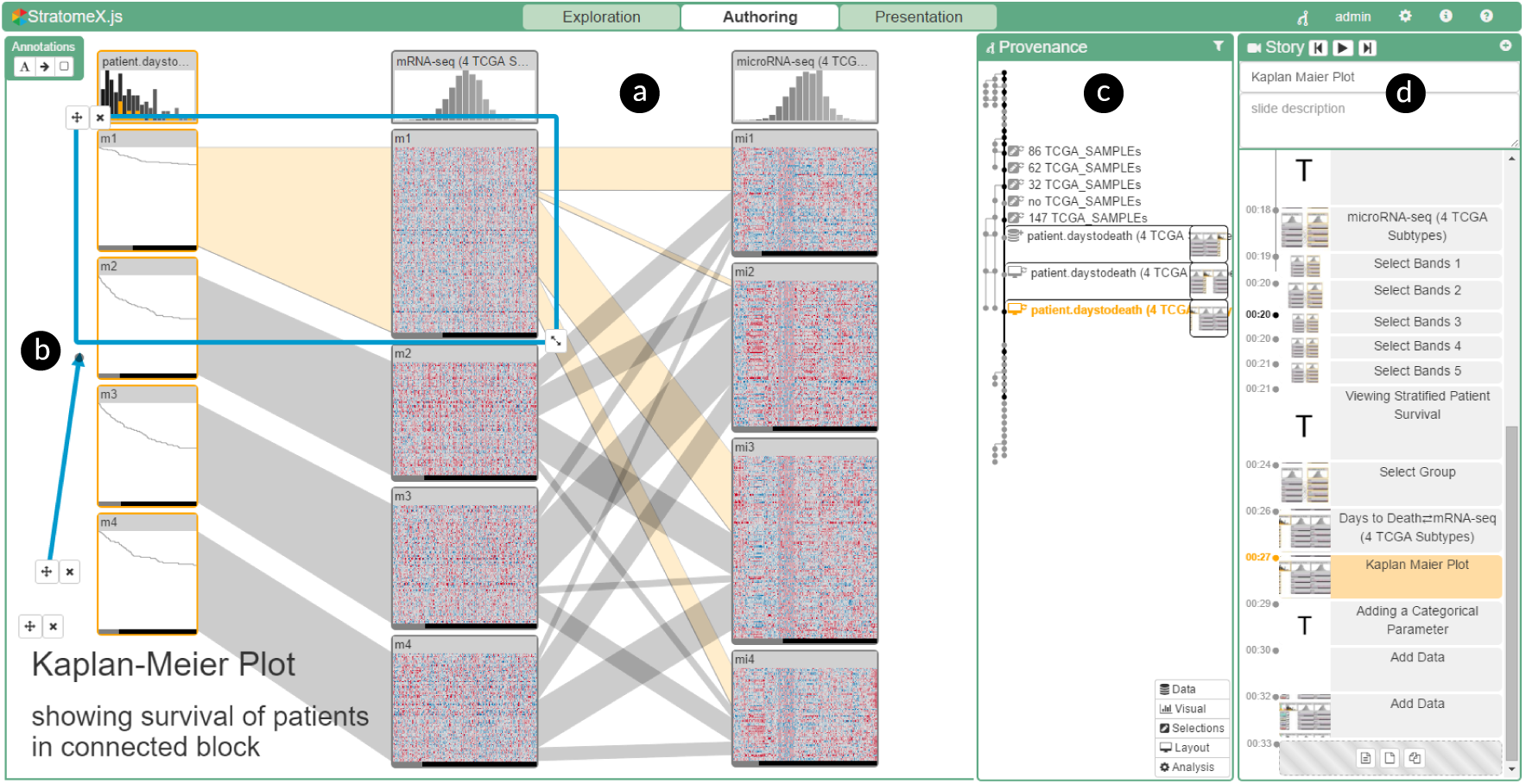
*Screenshot of CLUE applied to the StratomeX technique (a) in authoring mode. An annotation (b) highlights relevant aspects. The provenance graph view (c) and story view (d) show the history of the analysis and a Vistory being created*. *Vistory:* http://vistories.org/v/stratomex.

**Figure 2:**
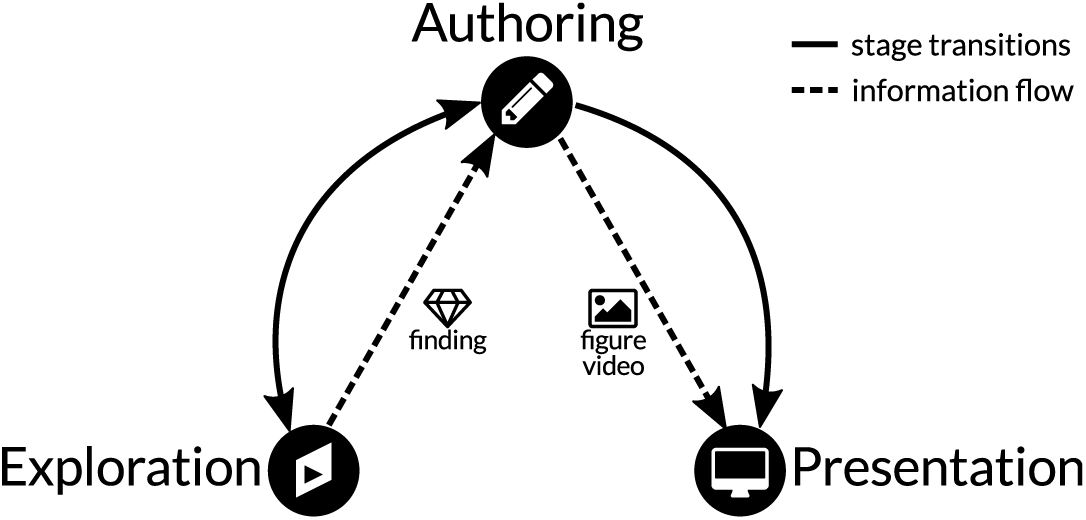
*Traditional workflow and information flow for visual data exploration and presentation of discoveries. Dashed edges indicate information flow, solid edges show transitions between stages. The information flow is sequential and different tools are used in each stage*.

In the exploration stage of a data-driven research project, analysts apply interactive visualization and analysis tools to gain new insights. Then they document the discovery process for reproducibility and presentation. Presentation can be in the form of text or figures in a paper or slide-deck, or in the form of interactive visual data stories. Visual data stories are increasingly popular as they are engaging and can communicate complex subject matters efficiently [LRIC15]. Only in rare cases, however, can static figures or visual data stories be created straight out of the exploratory tool. Instead, an authoring stage, in which artifacts, such as screenshots, are sorted, edited, and annotated is necessary. In the case of interactive data stories, this process often consists of custom development of software. The final product is subsequently used to present a finding to a consumer. A consumer in this context can be, for instance, a reader of a news article, a reviewer or editor of a scientific publication, or a colleague of the analyst, trying to understand a finding and the process of its discovery. This workflow corresponds to the storytelling process introduced by Lee et al. [LRIC15]. The authoring stage described here includes their “make a story” and “build presentation” stages, since the story being told is a scientific discovery not requiring a distinction between scripter (author) and editor.

In a visual exploration process, findings are often captured by taking one or multiple screenshots of the visualizations, or by creating a screen recording that shows the steps that led to the discovery. Static images, however, cannot tell the story of the visual discovery, as they cannot convey information about the exploration process. Videos are difficult to create, edit and update, and also do not capture the full analysis process. Furthermore, the tools used for exploration are in many cases not suitable for authoring and presentation. Neither images nor videos allow an exploration to be continued, and both prohibit users from asking additional questions. Given the sequential information flow and the separation of tools, it is inefficient for the analyst and creator of the story – and even impossible for the consumer – to work back from an artifact used for presentation to the exploration stage. The lack of a back-link from the curated story to the exploration stage and the underlying data makes it impossible (1) to reproduce and verify the findings explained in a figure or video and (2) to extend an exploration to make new discoveries. In this work, we propose a comprehensive set of solutions to these problems.

Our primary contribution is CLUE, a model for reproducing, annotating, and presenting visualization-driven data exploration based on automatically captured provenance data. We also introduce Vistories– interactive visual data stories that are based on the history of an exploratory analysis that can be used as an entry point to reproducing the original analysis and to launching new explorations. Figure 3 shows our proposed CLUE model. All information flow is routed through a central component, and all stages use the same universal tool. This allows the seamless stage transition indicated by the solid edges.

**Figure 3:**
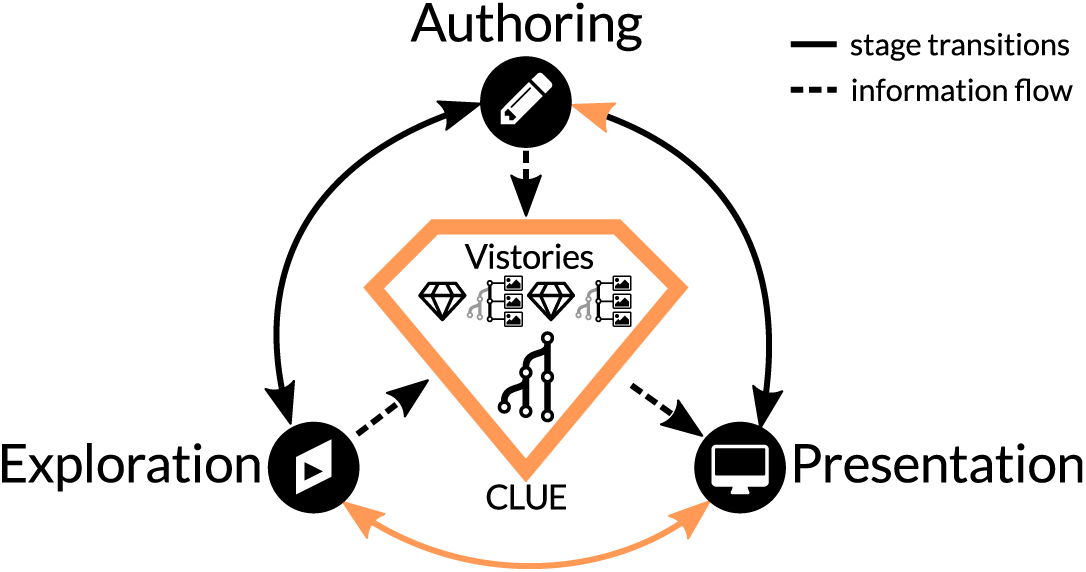
*Information flow and stage transitions using the CLUE model. The provenance graph of an exploratory session and Vistories (interactive visual stories) are in the center. Solid edges indicate possible stage transitions, dashed lines indicate information flow. In the exploration stage, provenance data is generated; in the authoring stage a Vistory is created by curating provenance data, which is then used in the presentation stage. Note that consumers of a Vistory can also switch to any other stage*.

As a secondary contribution we present a prototype implementation and a discussion of how to integrate CLUE with other visual exploration tools. To demonstrate the overall CLUE model, we describe a Gapminder-inspired usage scenario based on public health data. In a second usage scenario, we apply CLUE to a more complex multi-step visual exploration of cancer genomics data.

## 2 Related Work

CLUE closes the gap between exploration and presentation using provenance data. Since our model is independent of the exploratory visualization techniques employed, we limit our discussion of related work capturing and use of provenance data, presentation and communication in visualization, and visual storytelling techniques.

### 2.1 Provenance

In the context of our work, provenance of the state of a visual exploration process refers to all information and actions that lead to it. The provenance of the visual analysis as a whole is comprised of the provenance of all states that were visited during the exploration.

Ragan et al. [RESC16] have recently characterized provenance in visualization and data analysis according to the *type* and *purpose* of the collected provenance information. The type of provenance information includes data (i.e., the history of changes to the data), *visualization* (i.e., the history of changes to views), *interaction* (i.e., the interaction history), *insight* (i.e., the history of outcomes and findings), and *rationale*, the history of reasoning. Our prototype implementation currently captures all of this information. Data and interaction provenance are captured automatically, while insight and rationale are captured through the externalization of a thought process by the user. CLUE enables this, for instance via annotations and bookmarks.

Provenance information is used for several purposes [RESC16]: for *recalling* the current state during the analysis, for *recovering actions* (undo and redo), for *replicating* (reproducing) steps of the analysis, for *collaborative communication* (i.e., for sharing the analysis process with others), for *presenting* the analysis, and for meta-analysis (i.e., for reviewing the analysis process).

*VisTrails* [BCS*05], for example, collects and visualizes provenance data for computational and visualization workflows for large-scale scientific data analysis. Users of VisTrails interactively model a workflow that produces, for instance, a visualization; the process of creating this workflow is tracked as provenance data. In CLUE the focus is not on modelling a specific artifact with a goal in mind but rather automatically capture the provenance of an interactive visual exploration process that may lead to discoveries that are later being told using integrated storytelling approaches.

The works by Heer et al. [HMSA08] and Kreuseler et al. [KNS04] discuss a concept for visual histories (i.e., provenance of a visual exploration) including types, operations, management, and aggregation and provide a prototypical implementation. However, in both cases, provenance data is used for exploration only and not to address storytelling aspects. Heer et al. pointed to storytelling based on provenance as future work.

Action recovery (undo, redo) is commonly integrated into software applications. Most tools, however, do not visualize the history of actions and rely on a linear undo-redo path, which makes recovery from analysis branches impossible. Exceptions to this implicit approach are integrated in *Small Multiples, Large Singles* [vdEvW13], *Stack’n’Flip* [SSL*12], and *GraphTrail* [DHRL*12]. In the former two, the history of the current artifact is explicitly shown at the bottom and implicitly through the strict left-to-right workflow. GraphTrail also supports branches in the history: It explicitly visualizes how a plot is derived from previous ones using basic data operations. However, only a fraction of the provenance of the visual exploration is being captured by focusing on the data operations leaving out the parameters of the visualizations, etc.

In regard to provenance, the paper by Shrinivasan and van Wijk [SW08] is most closely related to CLUE. The authors proposed a technique that integrates three views: Data, Navigation, and Knowledge. The data view contains the actual visualization of the data. The navigation view shows the exploration process in a tree (i.e., the provenance). Using the knowledge view, users can capture and relate annotations to document findings, assumptions, and hypotheses, and link them to specific states in their exploration for justification. The knowledge view in combination with the navigation view is then used to communicate findings. In contrast to their work, the CLUE model also covers aspects of storytelling: by enabling authoring, we allow users to produce concise and effective stories based on the original exploration. This linear narrative approach is closely related to the traditional workflow of publishing results with the additional benefit of having a back-link to the real exploration at all time.

### 2.2 Storytelling

Kosara and MacKinlay [KM13] highlighted the importance of visual data stories for visualization research. They defined a story as “an ordered sequence of steps, each of which can contain words, images, visualizations, video, or any combination thereof”. They further stated that “stories can thus serve as part of a finding’s provenance, similar to an event’s narrated history”. Different approaches can be used for telling a story. Stories are mostly told individually and linearly, but management of multiple stories and branching have also been proposed [LHV12]. The degree to which users can influence how the story is being told may vary from automated replay to crafting their own story.

In CLUE, we apply a narrative approach storytelling inspired by the work of Figueiras [Fig14a, Fig14b]. Figueiras discussed how to include narrative storytelling elements such as annotations and temporal structure in existing visualizations to enrich user involvement and understanding through story flow. Similarly, the authors of *VisJockey* [KSJ*14] noted that when integrating interactive visualizations into data-driven stories, user guidance how to interpret the visualizations are lacking by default. Therefore, the authors proposed the VisJockey technique that enables readers to easily access the author’s intended view through supplementing the visualization with highlights, annotation and animation. In *Tableau* [Tab16], storytelling features are integrated using an annotated stepper interface. This enables users to navigate through a series of interactive dashboards. These works have in common that they are dealing with existing visualizations and insights, purely focusing on the authoring and presentation state, yet neglecting the underlying process of how the insights were discovered.

*Ellipsis* [SH14] is a domain-specific language for creating visual data stories based on existing visualizations. Journalists can combine visualizations, define triggers, states, annotations, and transitions via a programming interface or a visual editor. This allows them to define a wide range of story types, including linear, non-linear, interactive, automatic, and stepped stories. However, Ellipsis is only concerned with creating scripted stories, and does not utilize the visual data exploration process that leads to an insight.

With regards to storytelling, the work most closely related to CLUE is by Wohlfart and Hauser [WH07]. Their technique allows users to record interactions with a volume visualization, modify this recording, and annotate it in order to tell a story. In their work, however, the recording or capturing has to be actively triggered, and only a linear story can be captured. When users press a record button, they typically already know what they want to show. CLUE, in contrast, captures all actions and exploration paths, allowing the user to extract or base their story on the provenance of the analysis.

Storytelling approaches prompting user involvement through play and pause techniques were explored by Pongnumkul et al. [PDL*11] to support navigation in step-by-step tutorials. Adobe Labs [Ado16] provides a Photoshop plugin to create such tutorials: users can first record actions and then author them.

In summary, existing storytelling tools focus on how to tell a story, but rarely base the story on provenance data. The CLUE model solves this by providing links between points in the story to corresponding states in the exploration. This allows users switch freely and easily between exploration and presentation.

## 3 CLUE Model

The CLUE model bridges the gap between exploration and presentation. Its backbone is a rich provenance graph that contains all actions performed during the exploration. This includes exploration paths that led to findings, but also the dead ends encountered by the analyst. By putting the provenance graph at the center of our model, we are able to break the strict sequential order of the exploration, authoring, and presentation stages (see Figure 2) that dominates traditional workflows. CLUE allows users to seamlessly switch between **Exploration Mode, Authoring Mode**, and **Presentation Mode**. This integrated and flexible process is illustrated in Figure 3, where solid lines indicate the possible transitions between stages, and dashed lines represent information flow.

Provenance data makes scientific findings more reproducible and can also provide the basis for authoring stories. In CLUE, users create stories by defining a path through the provenance graph. We call such a provenance-based story a ***Vistory***. States in the story can then be enriched with highlights, textual annotations, and, if desired, timed for automatic playback. Hence, the resulting story is not an artificial composition of visualizations, but a curated version of the actual exploration. Most importantly, this deep integration of the provenance graph introduces the back-link from presentation to exploration. Vistories can be shared and encourage collaborative visual data analysis. Consumers can step through a story, but also switch to the exploration mode and interactively build upon the previous analysis to gain new insights. Vistories also make the exploration process more efficient, as a user can revisit states and apply changes, without redoing all steps to reach a particular state. Therefore, Vistories are more than visual data stories as defined by Lee et al. [LRIC15], since they allow consumers to continue the visual exploration in-place and build new stories themselves.

Figure 4 illustrates three scenarios that show how users can switch between modes. In the first example (Figure 4a), the process starts by investigating the data in exploration mode. After several iterations, the analyst discovers a finding worth presenting and switches to authoring mode to create a Vistory. Finally, the analyst previews the Vistory in presentation mode and makes it available to others. In the second example (Figure 4b), an editor creates a Vistory in the authoring mode, starting with an existing analysis session. After previewing the story in presentation mode and continued editing in authoring mode, the editor notices that the content of the story could be improved, and switches back to exploration mode in order to refine the visualization. Subsequently, the user returns to authoring mode and finishes the Vistory. In the last example (Figure 4c), a consumer starts by watching an existing Vistory. In that process, the consumer becomes curious about the consequences of adding another dataset to the analysis. The consumer switches to exploration mode and picks the relevant state, from where she can start her own analysis. Based on this new analysis, she creates a new Vistory that can be shared with collaborators.

**Figure 4:**
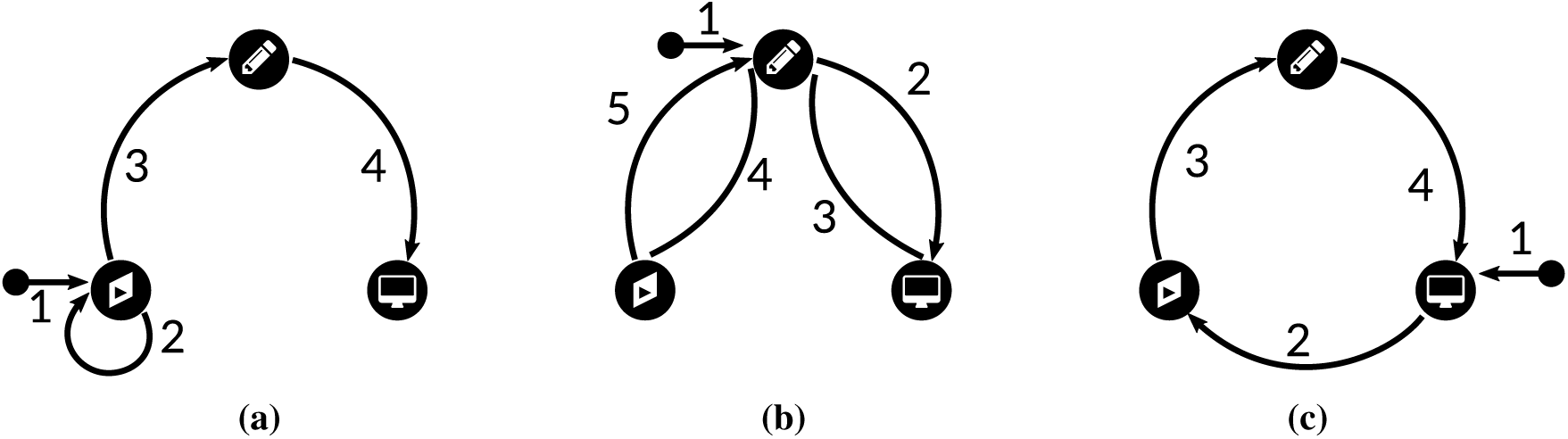
*Three examples of transitions within the CLUE model, highlighting different entry points. Numbers indicate the order in which the stages are visited by the user*.

These simple workflow examples illustrate that users can enter the process in any mode and switch freely between modes. Note that, from a conceptual point of view, switching from exploration directly to presentation is possible without going through an authoring step. While this can be useful when the only goal is to reproduce an analysis, authoring is required as a transition between exploration and presentation in practice to create an informative and concise story.

## 4 Realizing the CLUE Model

In this section we demonstrate the practical use of the CLUE model. We discuss how visual exploration tools can be extended by adding provenance capturing, authoring, and presentation capabilities. We also describe a prototype library for capturing and managing provenance and story data as well as a prototype that demonstrates the flexibility and efficacy of our model.

The library is used by a *Visual Exploration Application*. It is important to note that the Visual Exploration Application is not restricted to a specific visual exploration tool, but only has to comply with a set of basic requirements (see Section 5) and make appropriate calls to the library. The application is shown in all CLUE modes, although it is set to read-only during authoring and presentation.

The CLUE library consists of several building blocks. At its core is the provenance graph data model that forms the back end of CLUE that is used to store the captured exploration process. The other important components of the library are the provenance view and the story view.

The *Provenance View* provides a scalable visualization of the provenance graph. In exploration mode, a simplified version of the graph gives the analyst an overview of the provenance, by showing, for example, the states leading to the current one. When the system is in authoring mode, the provenance view is used for navigating and selecting the states of the exploration process that should be part of the story.

The *Story View* visualizes the elements of the story. Depending on the selected mode, the view shows different features. In presentation mode, it is essentially a stepper interface for the curated story. When the system is in authoring mode, it enables users to create, manage, and edit stories.

The visual components can be active in more than one mode and show different levels of detail depending on the mode. A switch between modes results in the addition, adaption, or removal of certain parts of the user interface. We apply animated transitions to support users in maintaining their mental model during mode changes. In addition, the current exploration state along with the mode used are encoded in the URL of the visual web application. This allows users to conveniently share states and Vistories by exchanging links.

### 4.1 Provenance Graph Data Model

The provenance graph data structure used in CLUE consists of four node types: *state, action, object*, and *slide*. Figure 5 illustrates the relationships between the different node types. An action transforms one state into another by creating, updating, or removing one or multiple *objects*. A state consists of all *objects* that are active at this point of the exploration. A *slide* points to a state along with annotations and descriptions explaining the state. A Vistory is made up of a sequence of slides. Switching between slides triggers actions that transition to the state associated with the target slide.

**Figure 5:**
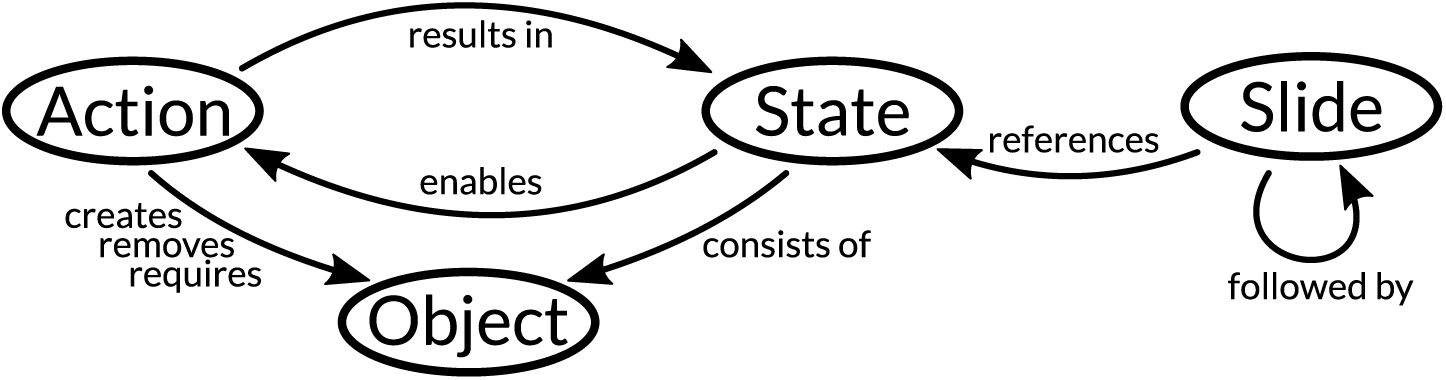
*The provenance graph data model consists of four different node types that are connected with each other by one or more edges*.

Object and action nodes are generic, and refer to the application-dependent implementation. In order to improve the visualization, additional metadata about objects and actions is stored. For objects we also store a *type*. We distinguish five types: data, visual, layout, logic, and selection. For actions we also store an *operation*: create, update, and remove.

Data actions deal with the addition, removal, or subsetting of datasets within the application. An example of a data action in our Gapminder usage scenario is the assignment of an attribute to an axis of the plot. Visual actions (e.g., switching an axis to logarithmic scale) manipulate the way datasets are shown to the user. Layout actions manipulate the layout of a visualization (e.g., manipulating the axes order in parallel coordinates or hiding the categorical color legend in a Gapminder plot). Logic operations, such as triggering a clustering algorithm, are concerned with the analytical aspect of the applications, Finally, selection actions encompass user-triggered selections of data in the visualization.

A state is characterized by the sequence of actions leading to it. Therefore, restoring a state is achieved by executing its corresponding actions. Jumping from one state to another is implemented by reverting the actions from one state to a common ancestor and executing the actions necessary to reach the target state.

In addition, for the purpose of transitions, the action sequence is compressed before being executed by removing redundant actions. A sequence of five selections, for example, will be replaced by the last one, since all the others are just intermediate states that do not influence the final selection. Similarly, when a create-and-remove action is associated with the same object, we remove both actions, as they neutralize each other. This compression avoids the execution of superfluous actions.

### 4.2 Provenance Visualization

As provenance graphs grow quickly during the exploration process, it is challenging to develop an effective visualization for it. Our provenance visualization is based on a node-link tree layout that we combine with a Degree-of-Interest (DoI) function to adapt the detail level of nodes [Fur86]. An example of the provenance visualization for one of our usage scenarios can be seen in Figure 1(c), and a close-up in Figure 6.

**Figure 6:**
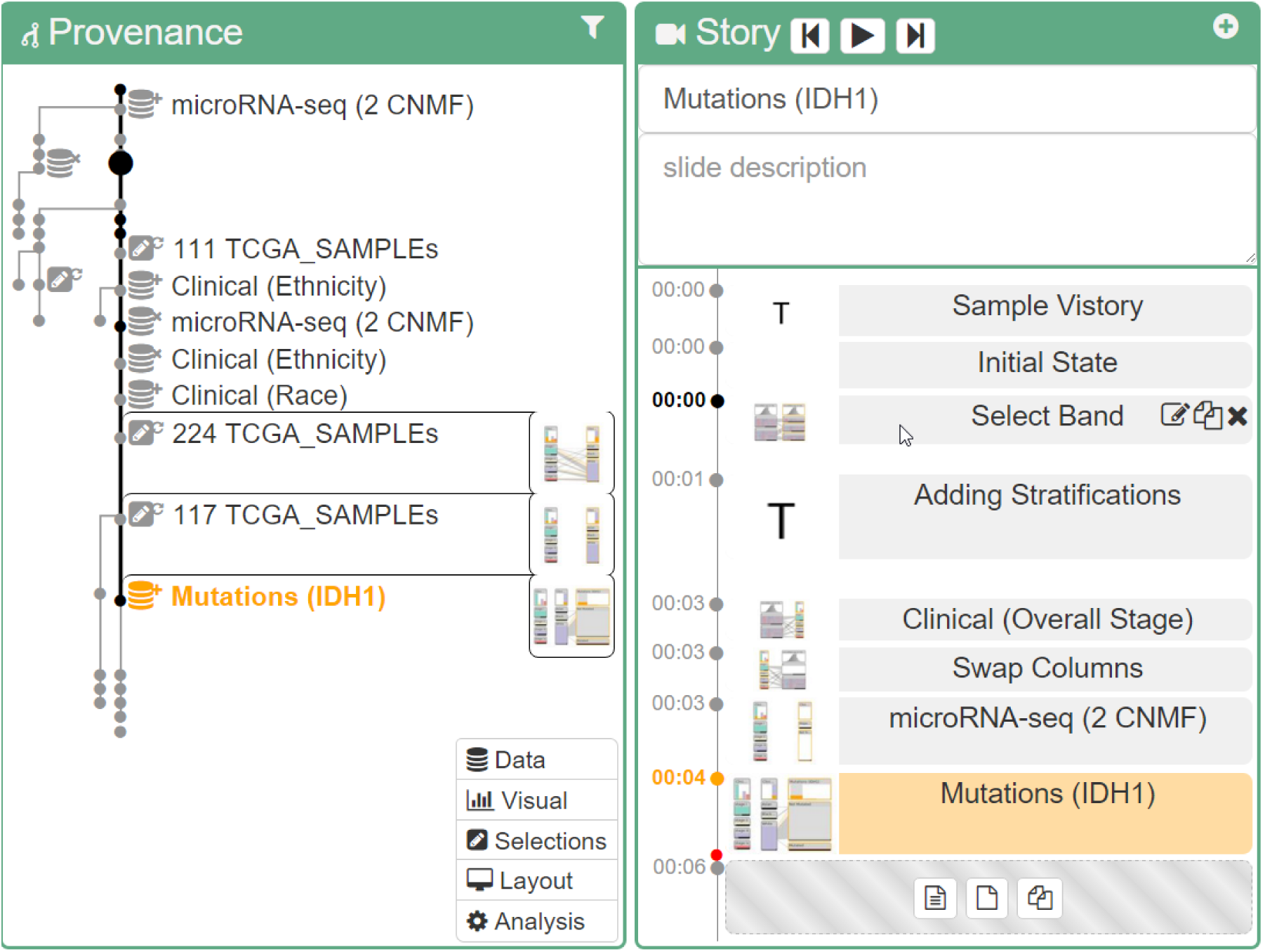
*Close-ups of the provenance and story views. Based on the DoI, exploration states are represented at different levels of detail in the provenance view. The structure of the Vistory, shown on the right, corresponds to a path through the provenance graph, which is shown as a thick black line in the provenance visualization. Both the active slide in the story view and the associated exploration state in the provenance graph are highlighted in orange*.

#### Layout

We use a vertical node-link layout as the basis for the provenance graph visualization. Nodes represent states, while edges represent actions transforming one state into another. However, instead of using a plain balanced tree, we reorder and skew it such that the currently selected node and all its ancestor nodes in the path to the root are aligned vertically on the right side. The remaining nodes and branches are then lined up on the left side. This strategy leaves space for details of the selected nodes and their ancestors, including labels describing the state and thumbnails previewing the state of the visualization. However, the layout needs to be updated when the user selects a different state. We use animated transitions to convey such changes.

#### Detail Level

We assign each node a DoI value that is influenced by several factors, including the current state selection, whether the state is an ancestor of the selected state (the distance term of the DoI function), and several filtering options that can be defined by the user (the a priori interest term of the DoI function). The DoI computed for each node is then used to adapt its representation with a combination of semantic and geometric zooming. We distinguish between four levels of increasing node detail.

**Level 1:** the state represented as a bullet

**Level 2:** icon describing the action associated with this state

**Level 3:** label describing the action associated with this state

**Level 4:** thumbnail of the application at the given state

Nodes of all detail levels are shown in Figure 6.

#### Interaction

Users can interact with the provenance graph visualization in several ways. Selecting a node will show the corresponding state in the application. At the same time the selection of a node triggers a re-layout of the provenance graph visualization, since the DoI values of the nodes change according to the selection.

The user can bookmark a state for later use, add tags, or add notes. Additional metadata can then be used to filter the states in authoring mode, which enables a more efficient story editing process.

To ensure reproducibility, it is important to be able to prevent a user from modifying the provenance graph. However, when authoring, the editor might want to improve a previous state, for example, by changing an axis scale from linear to logarithmic. Rather than allow the user to change existing states, we create a new branch. However, when starting a new branch, a user would need to redo all other actions that came after that was previously updated. We address this problem by allowing the user to apply a subbranch of a provenance graph to any other state. By dragging one node onto another, the actions are replayed automatically if possible. An example where this is not possible is when the user seeks to manipulate an object that has already been removed in one of the earlier states.

The consequence of preventing users from modifying the provenance information is that the graph grows rapidly. Although our provenance visualization supports multiple detail levels, the design and implementation of a truly scalable provenance visualization was not the main focus of CLUE and is therefore open for future research.

### 4.3 Story Editor

A story is composed of a sequence of slides. We distinguish between two slide types: *text slides*, which contain introductory text and captions, and *state slides*, which are associated with a specific state in the provenance graph. Both slide types can be annotated using multiple methods, such as styled text, scalable arrows, and freely adjustable boxes.

#### Layout

We use a vertical layout to present the slides, where we use the y-axis as a pseudo-timeline. The higher a slide, the longer it will be shown when automatically playing the story. Similarly, the more space between two slides, the longer the transition between them. Both transition and slide duration can be manipulated by dragging the top and bottom border lines of the slide, respectively. We chose a vertical layout because of (1) the alignment of the story with the provenance graph, and (2) the better readability of horizontal labels.

#### Interaction

Vistories can be created and edited in various ways. Editors can (1) start with a default text slide, which is useful if they already have an idea about the story they want to tell, (2) extract the currently selected state and all its ancestors, or (3) extract all bookmarked states.

Individual slides can be rearranged using drag-and-drop. In addition, dragging one state node in the story editor will wrap the state in a state slide when dropped, which allows a user to quickly create complex stories. Note that individual state slides need not be in sequential order in the provenance graph. The system automatically resolves the path between the states and plays all necessary actions, as discussed in Section 4.1. We indicate the currently active story in the provenance graph by connecting its states using a thick line. The provenance graph is also fully linked with the story editor: selecting a state slide in the story editor highlights the corresponding state in the provenance graph, and vice versa.

#### Annotations

Each slide can be enriched with annotations that are shown as an overlay on top of the visual exploration application. The library currently supports three different annotation types: text, arrow, and box. All of them are available in the movable annotation toolbar, which appear top left when a slide is selected in authoring mode. Figure 1(b) contains examples of all annotation types in blue. Annotations are positioned relative to anchor points in the visual exploration application. Anchor points represent important visual elements in the application such as data points of the scatterplot in Gapminder. By linking annotations to anchor points, their positions can be better adapted to layout changes due to different screen resolutions and aspect ratios.

## 5 Implementation

The CLUE library is a plugin of *Caleydo Web* [GGL*15], which is an open source, web-based visual analysis platform focused on biomedical data. Caleydo Web applications can use the library in order to provide them with CLUE capabilities. The source code is available at http://github.com/Caleydo/caleydo_clue. The Vistories for the usage scenarios and the video can be found at http://clue.caleydo.org.

CLUE and Caleydo Web are written in JavaScript and TypeScript on the client side and Python on the server side. The provenance graph is stored in a MongoDB [Mon16] database. Individual visualizations are implemented in D3 [BOH11]. We use the headless scriptable web browser PhantomJS [Hid16] to generate screenshots on the server by replaying actions of the provenance graph. This is a generic approach to replaying the Caleydo Web application enhanced with CLUE without the need for any customizations.

In order to use the library in a visual exploration application, is must use the *command design pattern* [GHJV95] for all recordable actions. These actions will then be captured in the provenance graph. We demonstrate the integration of two different applications—Gapminder and StratomeX— in our usage scenarios (see Section 6). In Gapminder, for example, recordable actions include choosing attributes for individual axes, switching scales of axes, and selecting years and countries. More sophisticated applications, such as StratomeX, support a larger set of actions.

## 6 Usage Scenarios

We demonstrate the utility of CLUE for a variety of applications in two usage scenarios. The first is inspired by Hans Rosling’s Gapminder http://www.gapminder.org. It illustrates the workflow of how users interact with and switch between different CLUE modes. The second scenario reproduces parts of our recent Nature Methods publication about StratomeX [SLG*14], a visualization technique for characterizing cancer subtypes. It highlights CLUE’s reproducibility support and its applicability to scientific analysis and storytelling. We provide links to the interactive Vistories for both usage scenarios below the respective figures.

### 6.1 Gapminder

A historian based in Europe is interested in assessing the interplay between wealth and health over the last 215 years. In particular, he would like to visualize changes in European countries to present his findings to a colleague in America. To explore health versus wealth, he first assigns income per person to the x-axis and life expectancy in years to the y-axis. The size a mark in the scatterplot corresponds to the size of the population of a country for the currently selected year. Contintens are color-coded; Europe is shown in Purple, America in red, Africa in blue, and Asia in brown. He applies a linear-to-log transformation to the wealth data and is ready to explore. Moving the slider on the timeline from 1800 to 2015, he observes an overall trend for Western countries: when wealth increased, people lived longer. The populations of most African countries, however, continue to have low GDPs and low life expectancies, as shown in Figure 7.

**Figure 7:**
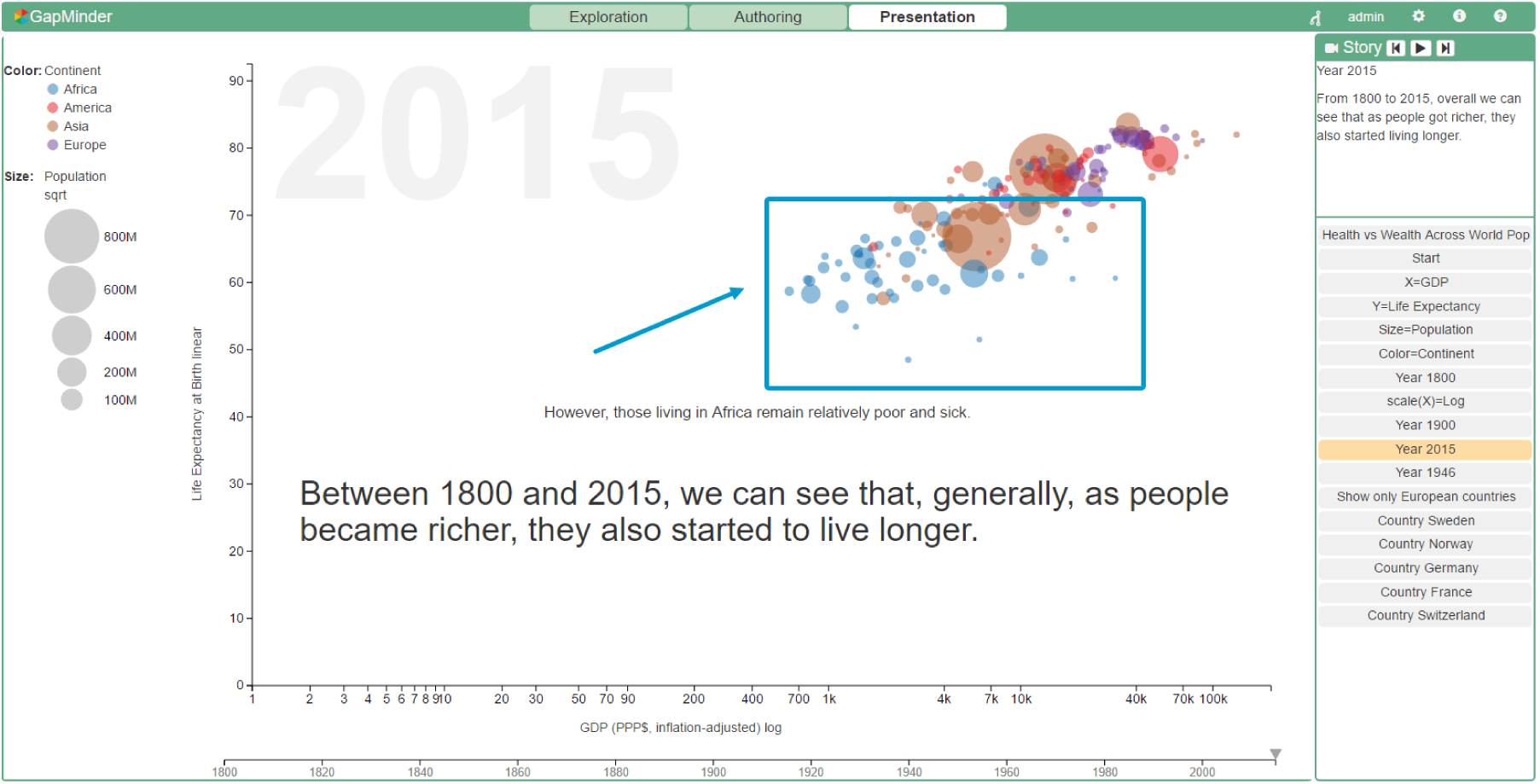
*Screenshot of a Gapminder-inspired application illustrating a Vistory in presentation mode. African countries are highlighted using annotations. **Vistory:*** http://vistories.org/v/gapminder.

To create a Vistory based on this finding, he switches to authoring mode, where he extracts all states captured in the provenance graph. In the story editor, he annotates the states using the text and arrow annotation tools and previews the story by clicking on the play control. Now, he would like to take a closer look at health and wealth in Europe in particular, so he switches back to exploration mode, where he uses the continent legend to select Europe from the chart. As a result, all countries in other continents are shown with reduced opacity. He goes to a pivotal year in European history: 1946, right after World War II. He evaluates the state of Europe regarding health and wealth, and extracts his findings to the story editor. In the story editor, he adjusts duration times and annotates key findings, moves to presentation mode to review, and finally shares a link to his Gapminder Vistory with his American colleague.

The American historian views the Vistory in presentation mode and decides that she would like to look at the state of Africa at the points chosen for Europe. To do so, she switches to exploration mode. Here she selects the node in the provenance graph where her European colleague initially selected Europe, and selects Africa instead. Her exploration is now captured on another branch of the provenance graph, and she is free to extract her own Vistory. To reproduce the analysis done for Europe for her subset, she applies the original subbranch to her new branch.

### 6.2 StratomeX

StratomeX is a visualization technique for cancer subtype characterization [LSS*12, SLG*14]. For this usage scenario, we ported the StratomeX visualization technique to Caleydo Web. In the following section, we describe how CLUE can be used to reproduce use cases described in the StratomeX paper [SLG*14]. To ensure data provenance, the authors provided the case study data. The exploration steps are illustrated in a supplementary video. However, the video shows only the final curated story and not the exploration as a whole.

We started to reproduce the case study by following the figure captions and the video step by step. This included frequent switching between exploration and authoring mode, while simultaneously creating the corresponding Vistory. In the end, we successfully reproduced Supplementary Figures 6(a) and 6(b) of the original paper. Figure 1 shows an intermediate screenshot of StratomeX in authoring mode, illustrating a central aspect of the technique. A similar picture is part of the original video. The left side shows StratomeX with annotations. On the right, the provenance graph of this analysis and the story view are shown.

With this CLUE-based implementation of StratomeX we were able to re-trace an analysis from a published paper and make this analysis reliably reproducible as a Vistory. This has all the benefits of the original research video, but was much easier and faster to produce. Moreover, consumers of the Vistory can go back to the analysis and, for example, check for confounding factors by adding other datasets, or start their own analysis to look for new findings. The Vistory is accessible at http://vistories.org/v/stratomex.

## 7 Discussion

### Separation of Concerns

The CLUE model contains three different modes: exploration, authoring, and presentation. Since each mode has a different focus, only the relevant information and visual elements relevant to the current mode are shown in the protoype implementation. The provenance graph visualization, for instance, is shown in a simplified version when in exploration mode, in full detail when in authoring mode, and not at all when in presentation mode. A different approach would be to show all views and elements at once and let the user decide which elements are useful for a specific task. Wohlfart and Hauser [WH07], for example, used such a unified interface. However, overwhelming the user with all possibilities can be distracting. In CLUE, we decided to reduce the elements in the interface by introducing three separate modes for exploration, presentation, and authoring, therefore making as much screen space as possible available for the data visualization.

### One Tool for the Whole Process

Lee et al. [LRIC15] raised the important question of whether one tool combining exploration, authoring, and presentation features is suitable and desirable. While Lee et al. state that it might be a promising endeavor, they also have concerns about it. The key question is whether a unified tool can cover all potential analysis needs. Creating a tool that allows all kinds of visual explorations is indeed challenging. We tackle this challenge by providing a library that existing visual exploration tools can use. However, we also consider adding options to import images, videos and websites into Vistories, so that the presentation can be complemented by the output of incompatible tools. For these parts of a vistory, however, we will not be able to provide a back-link to the analysis.

### Collaboration

Collaborative visual data exploration is a relevant and promising research direction. CLUE captures all actions performed during a visual exploration on a semantic level. Hence, extending this approach such that multiple users can perform an analysis based on the same provenance graph is a logical next step. In our current implementation, we support sequential collaboration, that is one user at a time can perform the analysis, but the user can change over time. However, our ultimate goal is to enable synchronous collaboration. This, will introduce several additional challenges, such as synchronization issues, or the visualization of such multi-user provenance graphs.

When introducing user management, we will also be able to restrict the operations allowed on a Vistory. For example, one could prohibit modification of a published Vistory and only allow a fork to be modified.

### Full Provenance

The current implementation of the CLUE model captures the steps carried out by the analyst during visual exploration. However, the datasets used during exploration, tools employed outside of our application, and which version of an application is used is not tracked. This information would be needed to truly reproduce every state of the exploration. The versioning of datasets, tools, and applications, however, is a subject of active research in other fields. For example, source code management systems such as Git and Subversion work well for all kinds of text files. In the biomedical domain, platforms such as Refinery [GPS*14] capture the provenance for the execution of workflows with all input and output data. Approaches based on Virtual Machines and Docker [Doc16] can be used to capture application versions. In the future, we plan to integrate all versioning approaches mentioned into a comprehensive solution that then allows full provenance tracking for data-driven visual exploration.

### Meta Provenance

CLUE currently captures only the visual exploration itself. All actions performed by the user in the authoring and presentation mode are not tracked. As users can jump to different branches of a provenance graph during the analysis, the sequence of actions performed by the user can only be reliably reconstructed via the timestamps of actions. An interesting future research direction is therefore to track the provenance of how the provenance graph was created. Capturing this meta-provenance graph would allow us to analyze the process of visual exploration and also the evolution and use of the CLUE model, including how stories evolve and how users collaborate.

### Scalability

Since provenance graphs grow very quickly, scalability is an inherent problem. To mitigate the scalability issue, the provenance graph visualization represents only states as nodes and actions as edges between them while hiding object nodes. We found this to be intuitive, since users tend to think in states. We found our DoI approach to be useful for managing medium-sized provenance graphs, but we realize that for large provenance graphs of complex analysis sessions, additional methods will need to be developed. One aspect we plan to investigate are user-specified states of interest that can influence both the visualization of provenance graphs and the selection of states for stories.

### State Selection

An important question when creating a story is which intermediate states to select given a target state. A simple approach is to use the path from the start of the exploration to the desired target state. This, however, may contain superfluous states and leads to long stories. Automatically identifying key states is challenging. While there are certain measures one could take, such as removing intermediate selections that were not pursued further, these assumptions are invalid in the general case. Manual annotations or bookmarks of states during an analysis are strong indicators for the relevance of states. These can be leveraged by suggesting key states for authoring and for emphasizing them in the provenance graph. We plan to investigate methods for encouraging users to externalize their assumptions and reasoning, which is also important with respect to reproducibility.

### Animated Transitions

Animated transitions are effective for communicating changes between different states. However, the CLUE library is independent of the visual exploration tool used. Therefore, a story can only suggest the duration of an animation between states, but the actual application must decide how the timing options are interpreted. Moreover, moving from one state to another can involve a series of actions. Currently, they are executed sequentially, but some of them could be executed in parallel in order to speed up the transition. Detecting independent actions, how they can be executed in parallel, and whether this can improve user understanding remain open research questions.

## 8 Conclusion and Future Work

We have presented CLUE, a model for capturing, labeling, understanding, and explaining provenance information of data-driven visual explorations. Based on the collected provenance information and by tightly integrating both exploration and storytelling in a generic model, users are enabled to switch from exploration to storytelling and back again. We also introduced a library implementing this model and two visualization tools that make use of this library.

As part of future work, we plan to perform meta-analysis on the recorded provenance graphs. This can be done on a single graph, (e.g., by detecting cycles and similar states) or on a collection (e.g., detecting common analysis patterns and action sequences). Both can be used to support users by pointing to possible next actions in an analysis or by indicating loops in the analysis.

In addition, we intend to launch a platform for sharing, viewing, and exploring Vistories along with the provenance graphs, data, and applications. Our vision is that an increasing number of visual exploration systems will capture the provenance graph of their analysis sessions and that these provenance and Vistory packages could be submitted along with a paper as supplementary material. This has the potential to simplify the job of reviewers, ensure reproducibility of the findings, improve the communication of the findings, and ultimately speed up scientific progress.

## 9 Acknowledgements

This work was supported in part by the Austrian Science Fund (FWF P27975-NBL), the State of Upper Austria (FFG 851460), and the US National Institutes of Health (U01 CA198935).

